# Exploring aggregation genes in a *P. aeruginosa* chronic infection model

**DOI:** 10.1101/2024.06.10.598275

**Authors:** Alexa D. Gannon, Jenet Matlack, Sophie E. Darch

**Author notes:** Corresponding Author 12901 Bruce B Downs Blvd, MDC7, MDC 3024, Tampa FL, 33612.

## Abstract

Bacterial aggregates are observed in both natural and artificial environments. In the context of disease, aggregates have been isolated from both chronic and acute infections. *Pseudomonas aeruginosa* (*Pa*) aggregates contribute significantly to chronic infections, particularly in the lungs of people with cystic fibrosis (CF). Unlike the large biofilm structures observed *in vitro, Pa* in CF sputum forms smaller aggregates (∼10-1000 cells), and the mechanisms behind their formation remain underexplored. This study aims to identify genes essential and unique to *Pa* aggregate formation in a synthetic CF sputum media (SCFM2). We cultured *Pa* strain PAO1 in SCFM2 and LB, both with and without mucin, and used RNA sequencing (RNA-seq) to identify differentially expressed genes. The presence of mucin revealed hundreds of significantly differentially expressed (DE) genes, predominantly downregulated, with 40% encoding hypothetical proteins unique to aggregates. Using high-resolution microscopy, we assessed the ability of mutants to form aggregates and identified 13 that were unable to form WT aggregates. Notably, no mutant exhibited a completely planktonic phenotype. Instead, we identified multiple spatial phenotypes described as ‘normal,’ ‘entropic,’ or ‘impaired.’ Entropic mutants displayed tightly packed, raft-like structures, while impaired mutants had loosely packed cells. Predictive modeling linked the prioritized genes to metabolic shifts, iron acquisition, surface modification, and quorum sensing. Co-culture experiments with wild-type PAO1 revealed further spatial heterogeneity and the ability to ‘rescue’ some mutant phenotypes, suggesting cooperative interactions during growth. This study enhances our understanding of *Pa* aggregate biology, specifically the genes and pathways unique to aggregation in CF-like environments. Importantly, it provides insights for developing therapeutic strategies targeting aggregate-specific pathways.

**Importance:** This study identifies genes essential for the formation of *Pseudomonas aeruginosa* (*Pa*) aggregates in cystic fibrosis (CF) sputum, filling a critical gap in understanding their specific biology. Using a synthetic CF sputum model (SCFM2) and RNA sequencing, 13 key genes were identified, whose disruption led to distinct spatial phenotypes observed through high-resolution microscopy. The addition of wild-type cells either rescued the mutant phenotype or increased spatial heterogeneity, suggesting cooperative interactions are involved in aggregate formation. This research advances our knowledge of *Pa* aggregate biology, particularly the unique genes and pathways involved in CF-like environments, offering valuable insights for developing targeted therapeutic strategies against aggregate-specific pathways.

## Introduction

Bacterial aggregates are observed in both natural and artificial environments. In the context of disease, aggregates have been isolated from both chronic and acute infections and can be formed by bacteria, archaea, and fungi (1–4). *Pseudomonas aeruginosa* (*Pa*) is one such bacteria. As an opportunistic pathogen, *Pa* causes disease in those whose immune systems or barrier functions are compromised. This includes those with chronic and acute wounds, medical devices, and chronic infection in the lungs of people with the genetic disease cystic fibrosis (CF). Once chronic *Pa* colonization is established, a large proportion of the infecting bacteria grow within airway sputum as aggregates (∼10-1000 cells) (1). In contrast, *in vitro Pa* growth results in the formation of large structures containing millions of cells(5, 6). Previous studies of *Pa* cells in large, well-mixed flask cultures, (macro-scale biofilm structures) have contributed significantly to our understanding of *Pa* growth, communication systems, and the mechanisms *Pa* utilizes to become tolerant to many antibiotics (7). However, growth in this context does not closely recapitulate that of actual infection – specifically the presence of aggregates. This divergence gives rise to a fundamental question in biofilm microbiology – how does spatially structured growth as aggregates influence infection?

While both biofilms and aggregates exhibit clinical resistance to antimicrobial agents, it is probable that the underlying mechanisms contributing to this resistance differ. As these phenotypes intersect, distinct differences between aggregates and biofilms have emerged. Notably, the exopolysaccharides Pel and Psl have been identified as crucial components for maintaining the integrity of *Pa* biofilms (8). However, even in the absence of these polysaccharides, *Pa* aggregates can still form in synthetic CF sputum media (SCMF2) (1). While the presence of Pel and Psl undeniably contributes to the tolerance of aggregates to therapeutic interventions like antibiotics and bacteriophages, the physical occurrence of aggregation appears to be closely linked to enhanced survival (1). Notably, the regulation of quorum sensing (QS) in *Pa* biofilms has predominantly been characterized as a binary on/off system for coordination of group behaviors and the production of public goods (9). However, our previous research has revealed that in aggregates, the response to QS signals is considerably more diverse (10). For instance, aggregates in the path of a signal may not uniformly respond to it, and alterations in the expression of the signal receptor (LasR) can partially counteract this variability. This stark difference between biofilms and aggregates is noteworthy, especially when considering that *Pa* utilizes QS to regulate approximately 300 virulence genes. There are also now several examples from the microbiome that demonstrate how microbes can manipulate the spatial organization of their population or community, inferring changes in virulence. The range of functional outcomes that are mediated this way include regulation of QS, increased antibiotic tolerance and cross-feeding of metabolites (10–13). This suggests that other virulent behaviors may be heterogeneously employed across individual aggregates during growth, and that the ability to modulate pathogen position can modulate pathogen virulence. This could have significant consequences for the evolution of bacterial populations and ultimately how we should approach them therapeutically.

These observed functional differences between aggregates and biofilms highlight an urgent need to understand the mechanisms *Pa* utilizes to regulate aggregate formation to develop new therapeutic strategies. In this study, we identify a subset of genes that play an integral role in aggregate formation in a synthetic CF sputum media (SCFM2). Many of these genes encode hypothetical proteins. Using high-resolution microscopy, we have uncovered a spectrum of spatial phenotypes when aggregate genes are disrupted. Using available omics tools, we have been able to predict groups of functional pathways that contribute to such variations in the spatial structure of developing *Pa* aggregates. By mixing aggregate mutants with the WT PAO1, we found that the presence of fully functional cells incites further spatial heterogeneity, suggesting that multiple finely tuned biological systems are required for successful aggregation. These data present the use SCFM2 as a tool to dissect the mechanisms *Pa* uses to form infection relevant aggregates. More specifically, we demonstrate how we can utilize this knowledge to pair micron-scale spatial structure with the physiological response of *Pa* cells within aggregates.

## Results

### A unique subset of genes is critical for *P. aeruginosa* aggregate formation

We have previously shown that *Pa* growth as aggregates occurs in distinct phases, where single planktonic cells form aggregates which undergo dispersal and form new aggregates (10). Although this can be observed *in vitro*, we still do not understand the mechanisms required specifically for aggregate formation. The goal of this study was to determine if a distinct sub-set of genes is required for *Pa* aggregate formation in a synthetic CF sputum media (SCFM2) compared to larger biofilm models. While SCFM2 replicates both physical and nutritional aspects of CF sputum, mucin is the only known required component for aggregation in this chronic infection model. *Pa* cultured in the absence of mucin is unable to form aggregates (1). We leveraged this mucin dependency to identify genes important for aggregate formation.

PAO1 was cultured in both SCFM2 and LB in the presence and absence of mucin for 8 hours. RNA sequencing (RNA-seq) revealed several genes that were significantly differentially expressed (DE) in the presence of mucin (Figure 1). Of these DE genes, 40% were identified as hypothetical proteins i.e. no known function. Only two genes were determined to be significantly upregulated in SCFM2 with mucin: PA1530 and PA1531. Referenced as hypothetical proteins in *Pa*, orthologs in other *Pseudomonas spp*. also have no known function. Of the 17 significantly DE genes, 88% were downregulated in the presence of mucin. We found that 5 of these genes are ncRNAs with known associations with *Pa* biofilm regulation: *phrS, crcZ, rsmY, rsmZ*, and P30 (14–16). Within known *Pa* quorum sensing (QS) systems, only *pqsC* expression met our significance cutoffs. We found that the *pqsABCDE* operon is consistently downregulated at 8 hours in the presence of mucin (SCFM2 and LB), suggesting PQS signaling is less important once aggregates have formed (>4 hours).

**Figure 1.**
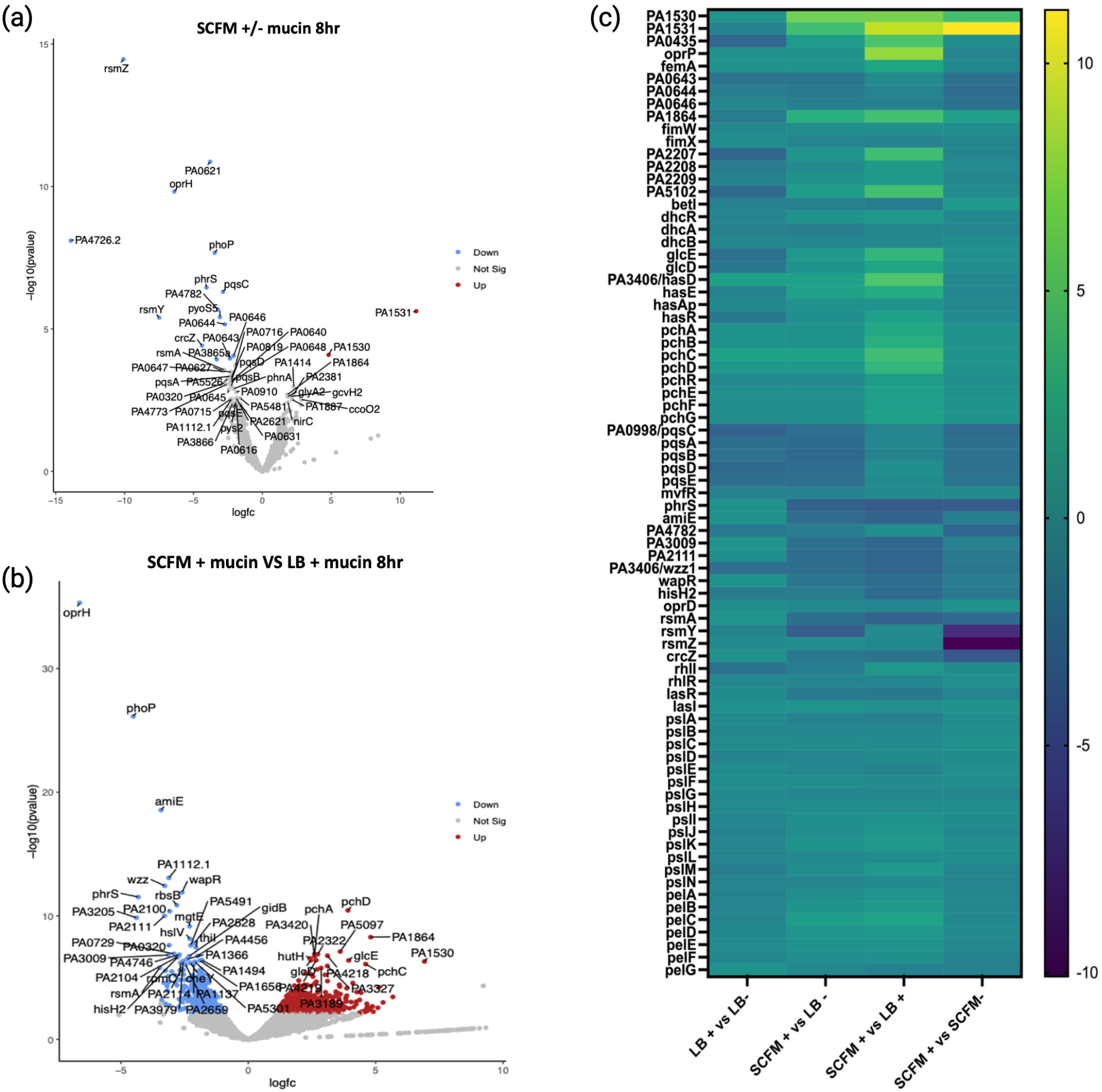
Differentially expressed genes in aggregates. (a) Volcano plot of differentially expressed genes in SCFM+ mucin compared to SCFM-mucin at 8 hours of growth. Upregulated genes are shown in red, downregulated genes are shown in blue, non-significant genes are shown in grey. 50 most significantly differentially expressed genes are labelled. 3 biological replicates, significance cutoffs 2-fold change, FDR 0.05 (b) Differentially expressed genes in SCFM+ mucin compared to LB+ mucin at 8 hours. (c) Heatmap of genes of interest and associated genes in their operon/pathway at 8 hours. Heatmap also include several classical biofilm-associated genes such as polysaccharides and QS genes that notably do not show differential expression in aggregates.

Next, we compared *Pa* cultured in SCFM2 with LB broth containing mucin. This allows us to identify genes that are specific to the nutritional environment of SCFM2 (Figure 1). Like our comparison of SCFM2 +/ mucin, a large percentage of DE genes were hypothetical proteins. Many pathway mapping programs (KEGG or GOseq) tend to exclude hypothetical proteins from analysis, making these tools inappropriate for our data. Therefore, we chose to focus on the 25 most upregulated and 25 most downregulated genes from this analysis (Table S1). Of this group, 40% of genes were identified as hypothetical proteins. This comparison also revealed several hundred significantly DE genes between the two growth environments, approximately 13% of the total PAO1 genome. More than half of the DE genes were upregulated (52%) in SCFM2 when compared to LB. This is a significant contrast to the number of upregulated genes (PA1530 and PA1531) attributed to mucin comparing SCFM2 alone. Notable downregulated genes specific to growth as aggregates in SCFM2 include the RNA-binding regulator *rsmA*, O-chain length regulator *wzz*, and virulence regulator *amiE* (17). We also observed an upregulation of several genes in the *pch, fptABCX*, and *glc* operons, encoding for the siderophore pyochelin, a pyochelin receptor, and a glycolate oxidase, respectively (18).

To further explore where our RNA-seq data overlaps with other *Pa* growth environments we compared significantly dysregulated genes in SCFM2 aggregates with those identified in data sets from PAO1 grown on a biofilm disk, within a flow cell and in *ex-vivo* CF sputum (Figure S1)(19, 20). We identified >500 unique genes to SCFM2 and 420 genes similarly dysregulated by PAO1 grown in CF sputum, including PA0621, PA0643, PA0644, PA0646, PA0985, PA0998, PA1530, PA1531, PA4782, PA5102, PA3406. These genes notably do not include those most commonly associated with the development of larger biofilms, such as the polysaccharide encoding genes *pel, psl*, or their related QS regulated *las* and *rhl* operons (Figure 1c)(21–24).

We also found that *Pa* growth as aggregates results in the downregulation of ncRNAs and associated repressors that impede biofilm formation (15, 16, 25). These data suggest two things 1) there are biological and metabolic processes that are distinct to *Pa* growth as aggregates (Figure 1c) and 2) that aggregate formation requires coordination across multiple biological pathways, including several that are still poorly defined.

### Aggregation mutants display a range of spatial phenotypes

Our initial goal was to validate our RNA-seq data and identify which genes, when disrupted, result in a non-aggregating, planktonic phenotype. To do this, we cultured single transposon mutants of the 50 significantly dysregulated genes in SCFM2 to assess how disruption of the gene impacted the ability of *Pa* to form aggregates (Figure 2). Cultures were imaged with confocal laser scanning microcopy (CLSM) over 15 hours and image analysis software was used to create 3D digital renderings of individual aggregates. Objects were filtered by size based on the dimensions of a single *Pa* cell; objects with volumes between 1-5μm^3^ were identified as single cells, and objects with volumes >5μm^3^ were identified as aggregates.

**Figure 2.**
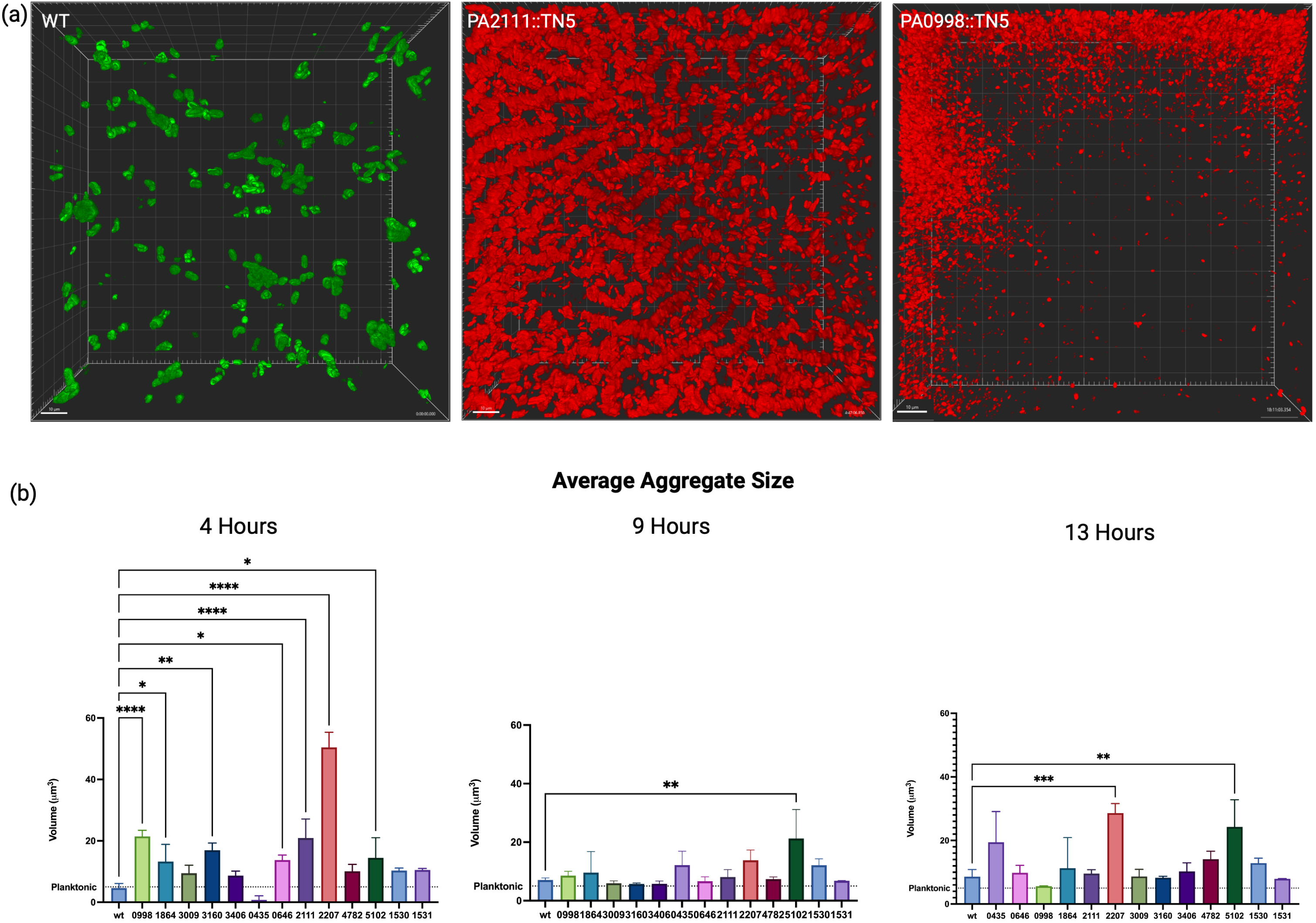
Transposon mutant aggregate phenotypes. (a) Examples of aggregation, with WT *Pa* shown in green and two representative mutants shown in red. Transposon mutant PA2111 is an example of the entropic phenotype, with long chains of stacked cells oriented along the coverslip. Transposon mutant PA0998 is an example of an impaired phenotype, with slow growth and loosely packed biofilm that becomes most apparent at later timepoints. (b) Average aggregate volume of transposon mutants over time. Mutant aggregates differ significantly from WT but vary over time. 3 biological replicates +/-SEM, significance calculated using one-way ANOVA (P value <0.0001) with Dunnett’s multiple comparisons test (alpha 0.05).

Objects with volumes less than 1μm^3^ were excluded from this analysis (as previously described(1)). Mutants were compared to WT aggregates quantitatively; characterized by the total biomass volume (μm^3^), and the average size of individual aggregates (μm^3^) over time as well as qualitatively, i.e. the presence or absence of aggregation (Figure 2, Figure S1).

These experiments identified 13 genes of interest, where *Pa* growth occurred as unique aggregation-deficit phenotypes when compared to WT PAO1 aggregation. We defined these aggregate phenotypes as either “normal’, “entropic” or “impaired” (Fig. 2a). It is important to note that the disruption of no gene resulted in a completely “non-aggregating” planktonic phenotype. Entropic aggregates are the result of *Pa* cells tightly packed side by side in long ‘raft-like’ structures. In comparison, impaired aggregates consist of loosely packed *Pa* cells with a number of planktonic cells during growth. A unique spatial arrangement, observation of the entropic phenotype in other studies of *Pa* is limited. Initially described, ‘entropic’ was used to describe *Pa* growth in relation to the physical forces or entropy present after the addition of polymers to media (26). A second example describes the significance of *Pa*’s O-antigen in mediating this spatial structure (26, 27).

We observed the entropic phenotype in cultures of Tn-mutants PA0435, PA0646, PA2111, PA2207, PA4782, and PA5102. Here, cells pack tightly side-by side, forming chains or stacks that elongate as the population matures. These structures form channels along the coverslip, overlaid by dense populations of disorganized planktonic cells and aggregates in the middle and upper layers of the Z-stack (Figure 2a). Entropic mutants were seen to maintain large planktonic populations, likely contributing to their accelerated growth rate compared to the WT. PA5102, PA2207, and PA0435 exhibited a loss of chain structures and corresponding population decline at 14 hours (Figure S1). Although we observe a general increase in growth rate compared to the WT, among entropic mutants this is variable. For example, PA0646 and PA2111 display a similar doubling time as the WT (80 minutes), while PA5102 and PA2207 exhibit robust, rapid growth that surpasses WT by 4 hours (94-108 minutes). PA0435 and PA4782 have an initial growth delay in SCFM2, entering exponential phase at 5 and 10 hours, respectively.

Additionally, both mutants form aggregates that are not significantly different from WT by volume (Figure 2b, Figure S1). PA5102 and PA2207 produce aggregates that are significantly larger (by volume) than WT, as well as maintaining their distinct morphology.

The impaired phenotype (characterized by loosely packed *Pa* cells) exhibits a similar growth rate to the WT PAO1, although significantly slower than that of entropic mutants (Figure 2a). This group includes the transposon mutants PA0998 (*pqsC*), PA1864, PA3009, PA3160 (*wzz1*), and PA3406 (*hasD*). All mutants except for PA1864 exhibit impaired or delayed growth, entering exponential growth around 9 hours as compared to 4 hours in WT cultures (Figure S1). The impaired mutant PA0998 (*pqsC*) demonstrates a significant growth deficit when compared to WT, revealing a previously unreported dependency on PQS quorum sensing for normal growth and wild-type aggregate formation in a CF lung like environment. PA0998 (*pqsC*) and PA3160 (*wzz1*) produce aggregates that are significantly larger than the WT at 4 hours, but transition to a primarily planktonic population of cells between 4 hours and the start of exponential growth at 9 hours (Figure 2b). The characteristic “impaired” architecture becomes most apparent between 10 and 15 hours of growth. PA1530 and PA1531 exhibit a “mixed” phenotype in which they exhibit impaired spatial structure during the first 10-12 hours of growth, after this time point, we observed the formation of entropic stacks.

### Predictive modeling reveals functional pathways specific to *Pa* aggregates

Next, we wanted to better understand the impact of disrupting aggregation genes on the physiology of *Pa*. Of the 13 prioritized genes (Table 1), 10 of those genes code for a hypothetical protein. We assigned predicted functions and pathways to each gene using our predictive modeling pipeline (Figure S2). This pipeline uses sequential and structural homology to assign gene functions and co-expression patterns to predict protein participation within known pathways. Interestingly, only three genes have known or suggested links to previously reported *Pa* biofilm functions: PA3406 (*hasD*) a component of the HasAD hemophore, PA0998 (*pqsC*) a component of the PQS quorum sensing (QS) pathway, and hypothetical protein PA1864 which has homology to a transcriptional repressor of FimWX mediated surface adhesion (Table 1) (28–31). PA1864 and *hasD* were upregulated in aggregate conditions 27.98 and 39.02-fold, respectively, while *pqsC* was downregulated -7.25-fold.

**Table 1.**
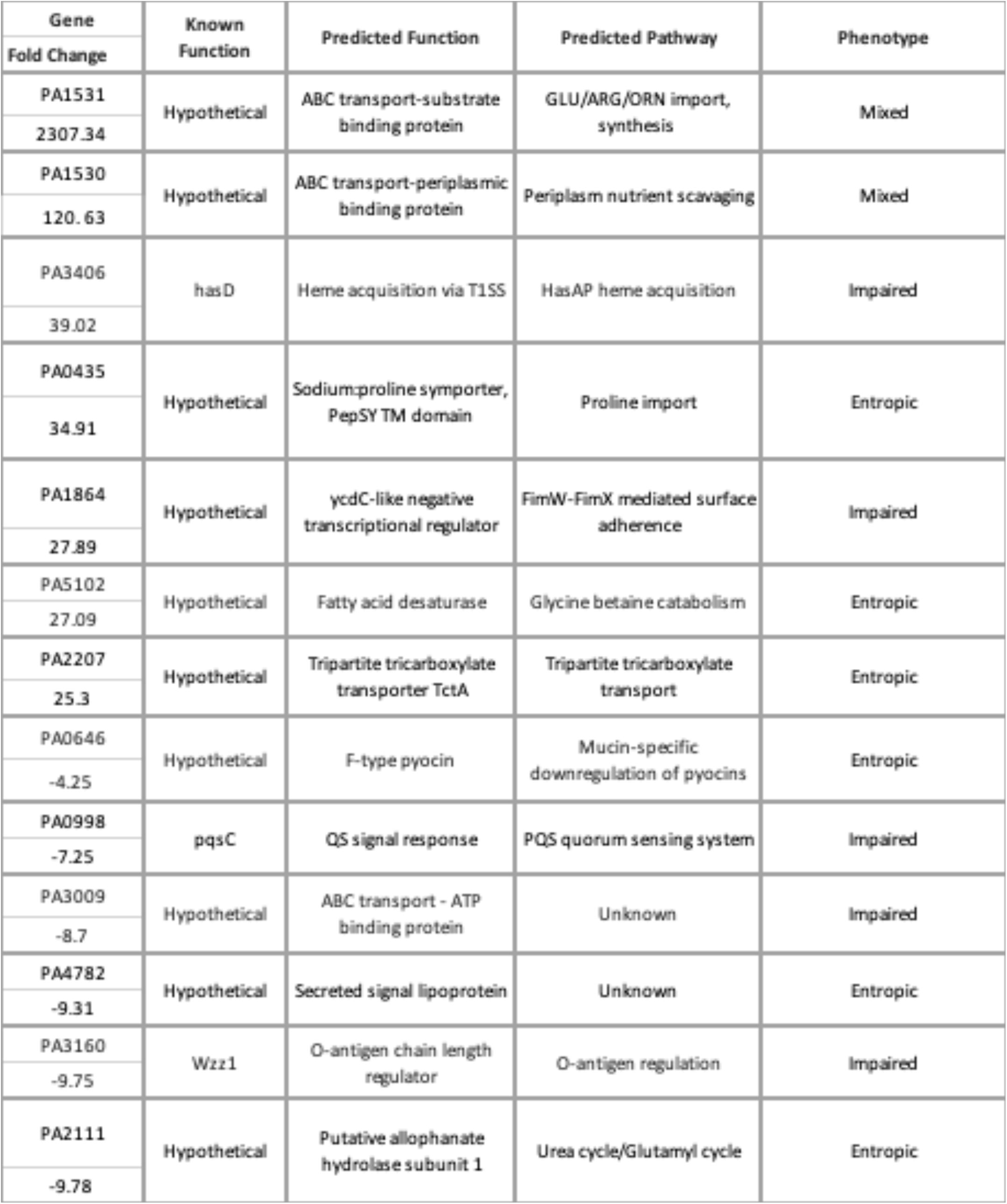
Genes identified as important for *Pa* aggregation. Table of genes, predicted function, pathway, and aggregate phenotype. Genes of interest identified from CLSM screening of transposon mutants. Gene function and predicted pathway for hypothetical proteins are informed from our prediction pipeline.

Using our pipeline, we are now able to propose a model of the genes and pathways critical for aggregate formation (Figure 3). We were able to group them by function, specifically by metabolism, iron acquisition, interaction and competition, surface modification, attachment, and QS related communication.

**Figure 3.**
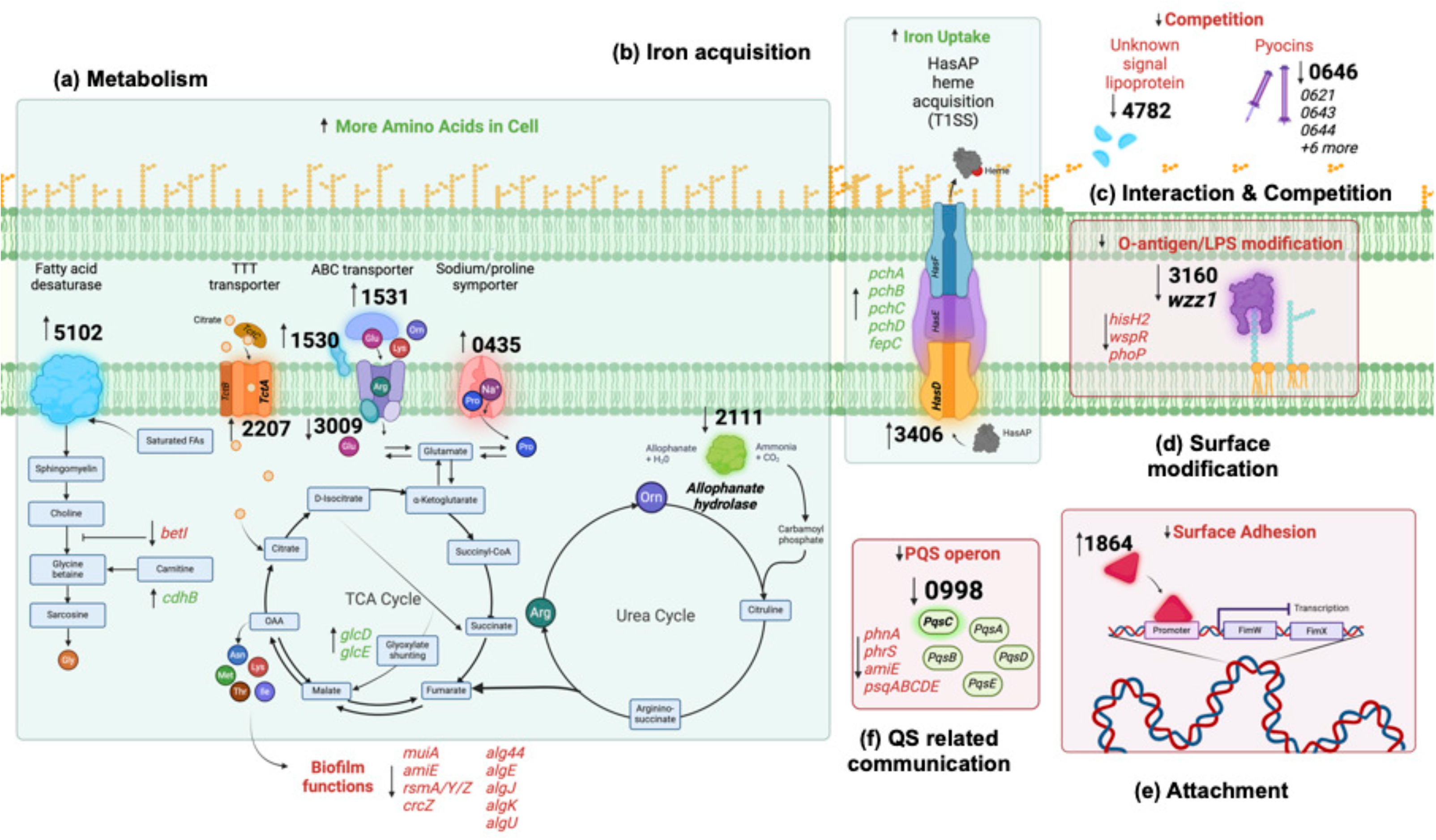
Proposed model of pathways involved in *Pa* aggregation. Hypothetical gene functions and pathways were predicted using computational pipeline. Genes of interest are in bold and grouped by function a-f. Upregulated functions are shown in green, downregulated functions are shown in red. Gene names within function indicate associated genes that were present in our dataset. Created with Biorender.

#### Metabolic shifts

Within the 13 genes, we identified an upregulation of multiple predicted amino acid transporters. PA1531 is predicted to be the substrate binding protein (SBP) of an ABC transporter and has high sequential and structural similarity to SBPs specific to glutamate, arginine, lysine, and ornithine. PA1531 is the most upregulated gene across all aggregate conditions, reaching a 2307.34-fold change in expression by 8 hours. PA1530 is predicted to be a periplasmic binding protein (PBP), which couples with specific ABC transporters in the periplasm to enhance nutrient uptake (32). PA1530 is also highly upregulated in aggregates at a 120.63-fold change. Our data strongly suggest co-expression and interaction between PA1530 and PA1531, indicating a critical role for amino acid transport and consumption during aggregate formation (Table 1). PA0435, a predicted proline/sodium symporter, is upregulated >30-fold. This protein also contains a pepSY domain, which is common in membrane proteins and has a proposed role in regulating local peptidase activity (33). PA2207 is upregulated 25.30-fold and is annotated in other *Pa* strains as *tctA*, the symport protein of the larger tripartite tricarboxylic transporter (TTT) assembly. TTT’s are not well characterized but have been shown to use ion gradients to bring citrate into the cell – although their substrate specificity is thought to be low (34, 35). In contrast, we observed a >8.70-fold downregulation of PA3009, which is predicted to be the ATP binding component of an ABC transporter.

Analysis of PA5102 and PA2111 provides support that distinct metabolic shifts occur as cells transition from a planktonic to aggregate growth style. Upregulated ∼27-fold at 8 hours, PA5102 is a predicted fatty acid desaturase involved in glycine betaine and carnitine catabolism, associated with the production of amino acids such as glycine (Figure 3). The utilization of this pathway is also supported by the downregulation of upstream inhibitor *betI* in aggregate samples at 4 hours, followed by the upregulation of both PA5102 and *dhcB*, which is involved in the conversion of carnitine to glycine betaine (36). PA2111 is predicted to be an allophanate hydrolase that is downregulated -9.78-fold in aggregates. Allophanate hydrolases have been shown to be involved in the conversion of allophanate and H_2_O into ammonia and CO_2_, ultimately leading to the formation of carbamoyl phosphate, a major input into the urea cycle.

This downregulation coupled with similar growth of PA2111 to the WT and a lack of other DE urea cycle-associated genes suggests that the urea cycle may not be utilized as heavily in aggregates as it is in planktonic cells (Figure S1).

#### Iron acquisition

PA3406 (*hasD*) is the inner membrane component of the HasAD hemophore and is upregulated in aggregates 39.02-fold at 8 hours, accompanied by upregulation of the remaining HasAP heme acquisition and *pch/fptABCX* pyochelin operons (Figure 1). Interrupting the HasAP secretion complex (and therefore, heme acquisition capacity) in a HasD transposon mutant leads to a growth delay and a loosely packed, impaired biofilm phenotype with a large planktonic population (Figure 2).

#### Interaction and competition

PA4782 (-9.31-fold change) is predicted to be a secreted lipoprotein with a signal peptide, although our pipeline was unable to assign a specific function or pathway due to lack of homology to proteins in other organisms. PA0646 is an F-type pyocin that is downregulated -4.25-fold and is part of a larger trend of mucin-specific pyocin downregulation (37)(Table S2). Pyocins are a secreted particle used by *Pa* to compete against other organisms and even other strains of *Pa*. We saw an interesting pattern of pyocin downregulation that is consistent and exclusive to mucin-containing samples and includes a mixture of R and F pyocins, with one S pyocin – pyoS5 (Table S2) (38). Transposon mutants of both PA4782 and PA0646 form entropic aggregates (Table 1).

#### Surface modification and adhesion

*Pa* as aggregates also exhibit a pattern of downregulation of well described extracellular components, three of which cause aggregation-deficient phenotypes. PA3160 (*wzz1*) is a regulator of O-antigen length on LPS and is downregulated 9.75-fold in aggregate conditions. These findings are accompanied by a trend of downregulation in other O-antigen and LPS modifying genes including *hisH2, wapR*, and *phoP*, suggesting a decrease in O-antigen and LPS modification in aggregates. More specifically, these data suggest that cells capable of forming aggregates adopt a specific outer membrane composition and undergo a distinct shift away from the production of some secreted factors. In comparison, temporal expression of these genes is critical for WT aggregate formation and growth.

#### QS related communication

Our dataset only identified significant differential expression of one QS system, when *Pa* exists as aggregates - PQS. PA0998 (*pqsC*) is significantly downregulated -7.25-fold in aggregates at 8 hours. The remaining PQS operon is also downregulated, although at lesser levels (Figure 1). Despite low expression levels at all timepoints, transposon mutant of *pqsC* shows a severe growth deficiency in SCFM2. Our data also indicates that tight regulation of the PQS quorum sensing system is integral for aggregate formation. *pqsABCD* are needed for synthesis of the PQS signal molecules (39). In the disruption of *pqsC, Pa* cannot form WT aggregates in a CF-like environment – timelapse microscopy reveals sparse populations composed mostly of planktonic cells (Figure 2a). The PQS system has been documented as critical for several components of biofilm formation, including iron chelation and induction of the oxidative stress response (40, 41). Gene expression patterns in our data also point to the importance of these two functions in the formation and maintenance of aggregates.

### Co-culture with the WT PAO1 results in further spatial heterogeneity

We co-cultured aggregate mutants (expressing GFP) with the WT PAO1 (expressing mCherry) in a 1:1 ratio in SCFM2. Using high resolution microscopy, we observed developing populations to assess three things: 1) the ability to restore the WT aggregate phenotype 2) if any mutant can outcompete the WT and 3) if the addition of WT cells influences the spatial organization of mutants. We found that the growth of fully functional WT cells with aggregate mutants resulted in changes in total biomass, average aggregate volume, number of planktonic cells (Figure 4b), and observed spatial organization (Figure 4a).

**Figure 4.**
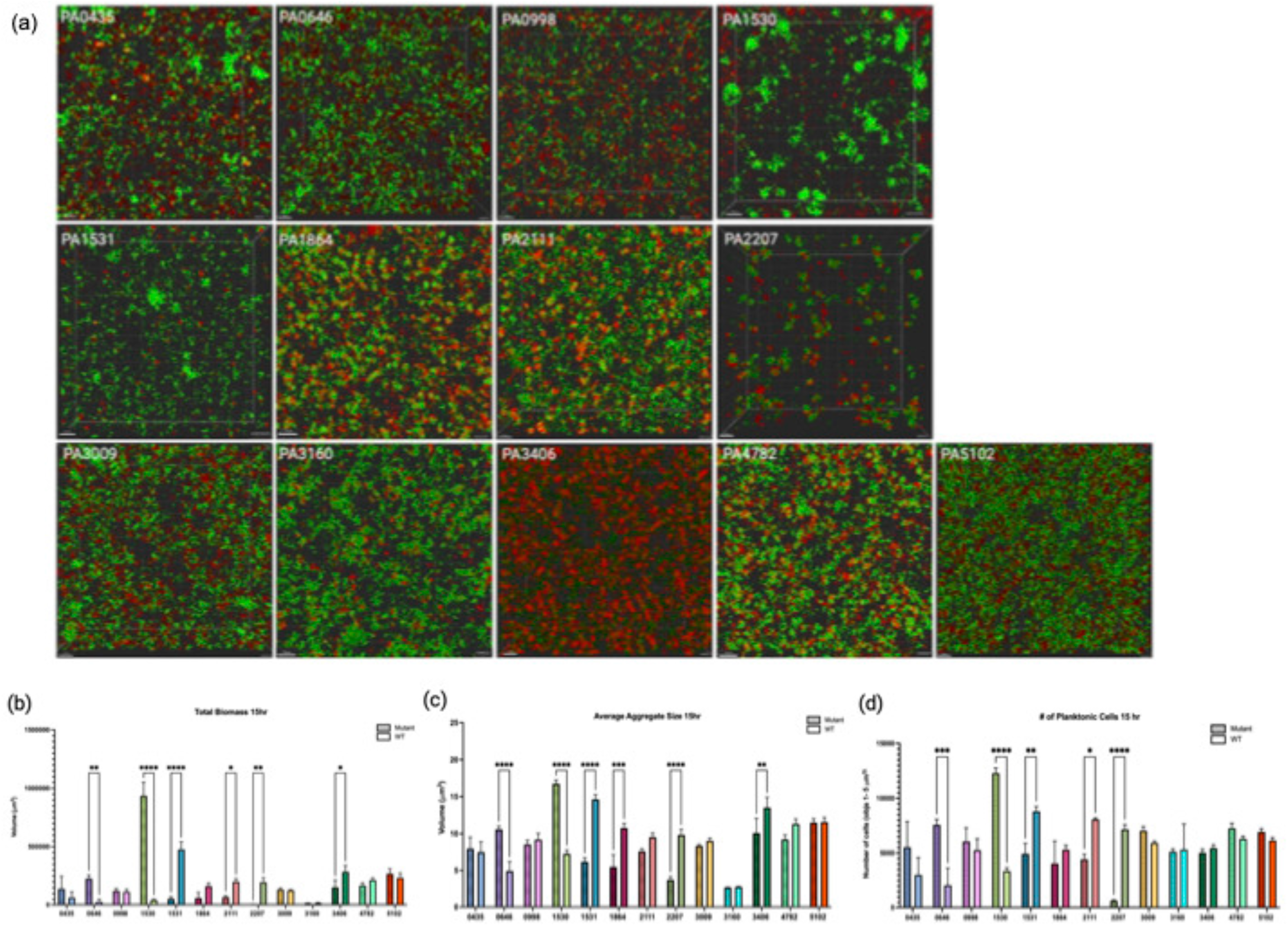
Co-culture of WT PAO1 and aggregation mutants. (a) confocal laser scanning microscopy (CLSM) of WT PAO1 (green, GFP) and transposon mutants (red, mCherry) co-cultured in SCFM2 at a 1:1 ratio after 5 hours growth. Scale bar is 10 μm (b) Comparisons of mutant and WT total biomass, (c) average aggregate size, and (d) number of planktonic cells at 15 hours. 3 biological replicates +/-SEM, significance of mutants vs. WT calculated using ordinary two-way ANOVA (P value <0.0001) with Fischer’s LSD multiple comparisons test (alpha 0.05).

We found that mutants PA4782, PA1864, PA0435, and 2207 restored ‘WT-like’ aggregation by volume when mixed with the WT. However, strains did not maintain separate clonal populations. Instead, we observed extensive mixing of strains. In contrast, PA0646 resulted in WT aggregate phenotypes, though strains maintained separate clonal populations. Strains did not form mixed structures and underwent separate dispersal events (Figure 4a). This is captured in PA0646, with higher overall biomass and aggregate volume at 15 hours, shortly after a WT dispersal event (Figure 4b). PA4782 showed no significant difference in any measure compared to the WT. PA1864 and PA0435 exhibited significant differences in both aggregate volume and the number of planktonic cells, with mutant strains producing smaller aggregates than WT.

PA3160 and PA3009 mixtures also revealed strains maintaining distinct clonal populations but did not restore WT sized aggregates. Although there were no significant differences between strains, these clonal populations formed extensive lawns of cells with distinct layers (Figure 4a). Finally, multiple combinations retained the original mutant phenotype. Specifically, PA1531, PA1530, PA 2111, PA5102, PA0998, and PA3406 mutants retained either an entropic or impaired aggregate structure. Interestingly, several mutants like PA3406 and PA5102 were able to incorporate WT cells into mixed entropic structures (Figure 4b).

From the opposite perspective, in many cases WT populations exhibited significant variation in total biomass, aggregate size, and planktonic populations between co-cultures (Figure S3). For example, when the PA1531 mutant is cultured alone, it forms significantly smaller aggregates, and has a reduced total biomass and planktonic population when compared to the WT. Conversely, when mixed, WT PAO1 exhibits significantly larger phenotypes when compared to its growth in the presence of other mutants (Figure S3). These findings strongly suggest that aggregation mutations can be ‘rescued’ in some cases. However, they also suggest that aggregation requires multiple biological processes, likely under precise temporal control.

## Discussion

Our hypothesis was that *Pa* aggregate formation requires a unique subset of genes when compared to much larger mm-scale biofilms. We have previously shown that the presence of mucin is a requirement for aggregation in our chronic *Pa* infection model of synthetic CF sputum (SCFM2) (1, 42, 43). In contrast, there is evidence that mucin promotes dispersal in *Pa* biofilms in an acute infection model, highlighting its potential importance in mediating the aggregate-planktonic lifestyle switch (44). Here, we leveraged this as a tool to control aggregation and study the differences between planktonic cells and aggregates of *Pa* in SCFM2.

### Polysaccharides are not required for early *Pa* aggregate formation

Our previous studies have identified the ability of *Pa* to form aggregates despite the absence of Pel, Psl or alginate (1). It is important to note that previous experiments provided qualitative data, simply examining the ability of PAO1 lacking polysaccharides to form aggregates. Here, using RNA-seq to compare planktonic cells with aggregates, we did not observe significant differential expression of the genes *pel, psl* or *alg* and their associated pathways. Neither did we see significant dysregulation within the associated QS pathways Las or Rhl. More specifically, there was no indication of these genes having a critical role in aggregation specific to any time point, mode of growth, or environment. Instead, our findings point to shifts in several metabolic pathways required for the switch from planktonic cells to aggregates in SCFM2. Many of these pathways have not been previously linked to aggregates and are not well defined, however they reveal novel insights into the metabolic needs of aggregates (Figure 1c).

### Aggregation requires temporal regulation of specific metabolic genes

Each of the 13 genes prioritized in this study demonstrate changes in expression between time periods of 2, 4, and 8 hours. Genes that are upregulated in aggregates at 8 hours, such as PA5102 and PA2207, may be required for adopting the slower metabolism and doubling time of aggregated cells. Mutants of these genes form entropic aggregates with rapid growth rates and large populations of planktonic cells. Disrupting genes that are downregulated after initial aggregation occurs (>4hours) also results in different mutant phenotypes, despite their perceived unimportance at this stage. This suggests a role in maintaining the aggregate once it has formed. Hypothetical proteins PA2111, PA3009 and PA4782, are upregulated at earlier time points. Mutants of these genes are unable to form WT aggregates, indicating early expression of these genes is required for aggregate formation. PA2111 is predicted to be a subunit of an allophanate hydrolase, part of a complex that converts allophanate into CO_2_ and ammonium, which eventually feeds into the urea cycle. Expression of this gene in SCFM2 cycles over time, peaking at 4 hours and decreasing to -9.78-fold downregulation at 8 hours (Figure 1c, Table 1). This is evidence of increased nitrogen levels, possibly attributed to amino acid catabolism during initial aggregate growth at 4 hours, creating urea as a waste product. This expression pattern is specific to aggregate growth in SCFM2, where amino acids are the primary carbon source. In LB, expression of PA2111 peaks later at 8 hours (Figure 1c). Studying gene expression patterns in this way provides new insight into aggregate metabolism and the impact of the microenvironment over time.

Amino acids are abundant in SCFM2 (and actual CF sputum) (45). Accordingly, several highly upregulated genes in aggregates (PA1530, PA1531, PA2207, and PA0435) form part of an amino acid importer (PA1530, PA1531, PA0435), or can feed directly into the TCA cycle (PA2207). PA1531 is consistently upregulated in SCFM2 +mucin samples, exhibiting > 2,000-fold upregulation compared to planktonic samples (Figure 1). Our computational pipeline predicts this hypothetical protein is the SBP of an ABC transporter, with specificity for arginine, lysine, ornithine, and glutamine, while PA0435 is a predicted proline/sodium symporter. We also see upregulation in SCFM +mucin of two probable amino acid aminotransferases (PA0870 and PA3139) that could facilitate this reaction (Figure 1c). PA2207 (TctA) is the transmembrane symport protein of a tripartite tricarboxylic transporter (TTT), which are not well characterized but have been shown to facilitate citrate uptake in several organisms (Figure 3). In parallel, PA5102 is a predicted fatty acid desaturase involved in glycine betaine metabolism and is upregulated in aggregates at 8 hours. This is preceded by the downregulation of repressor *BetI* at 4 hours, which acts as an inhibitor in the conversion of saturated fatty acids to glycine betaine (36). We also observe a cognate upregulation of *DhcABR*, which aligns with STRING co-expression predictions and is required in the final steps of the overall conversion of carnitine to glycine (Figure 1c, Figure 3, Figure S2). PA2207, PA0435, and PA5102 are entropic mutants with prolific replication rates (Figure 1). Evidence that repression of these highly expressed genes results in robust growth tells us several things about aggregate metabolism and its links to aggregate formation. First, the difference in growth rate between entropic and impaired mutants suggests that these spatial phenotypes are metabolically diverse (Figure S1). Second, that these mutations are not lethal suggests that *Pa* is able to compensate for their lack of function. Redundancy in the genome could explain this, where there are several homologs that could be co-opted to perform some of these functions.

### Some QS systems are redundant during aggregation

While the Las and Rhl QS systems are not significantly DE across our samples, our data suggests that tight regulation of the PQS QS system is integral for aggregate formation. PA0998 (PqsC) is significantly downregulated by more than 7-fold when compared to planktonic cells at 8 hours (Figure 1). Despite low expression levels at all timepoints, the transposon mutant *pqsC* has a severe growth deficiency in SCFM2. PqsABCD are required for synthesis of the PQS signal molecules (39). Here, we show that in the absence of PqsC, *Pa* cannot form WT aggregates in a CF-like environment, timelapse microscopy reveals sparse populations composed mostly of planktonic cells (Figure 2). To our knowledge, this is the first evidence of an explicit link between the ability of *Pa* to produce PQS and aggregate formation. The PQS system is well described as important for several components of biofilm formation, including iron chelation and the induction of oxidative stress response (40, 41). Gene expression patterns in our data also point to the importance of these two functions in the formation and maintenance of aggregates. We observed ∼40-fold upregulation of PA3406 (*hasD*) in aggregates at 8 hours, accompanied by upregulation of the rest of the *HasAP* heme acquisition and *Pch* pyochelin operons (Figure 1). Interrupting the HasAP secretion complex (and therefore heme acquisition capacity) in *hasD* transposon mutants leads to a growth delay and a loosely packed, impaired biofilm phenotype with a large planktonic population (Figure 2). Circumventing several steps in the TCA cycle, Glyoxylate shunting allows for the conversion of isocitrate to malate and is often observed in *Pa* isolates from human infections. Here in SCFM2 aggregates, we see an upregulation of glyoxylate oxidase subunits *glcDE* (46) (Figure 1, Figure 3).

One of our most interesting findings was that instead of an expected binary phenotype of aggregation or no aggregation, we identified several intermediate spatial structures between planktonic growth and WT aggregates. This suggests that failure to meet any one of the required pathways has the potential to inhibit WT aggregation to varying degrees. Approaching the study of aggregate spatial structure as a continuum instead of binary provides a platform to study how changes in the microenvironment shape spatial structure. Due to the nature of *Pa’s* sociality, it is likely that spatial organization within aggregates can impact the ability to share public goods, the use of gradient-driven communication, and antibiotic resistance. For example, the loose packing in impaired aggregates would likely cause public goods and signal molecules to have to travel farther, possibly exceeding their effective distance (10). The tight stacking in entropic mutants leaves little, if any, space between cells, reducing the distance that public goods and signals must diffuse to be shared (Figure 2). As a result, interactions that result from direct contact are more likely.

We decided to explore this further, by mixing fully functional WT cells with aggregate mutants. We asked if the spatial organization of mutants could be reverted to WT when cultured in SCFM2. Combinations of strains resulted in further shifts in spatial organization. For example, entropic aggregators PA4782 and PA2111 (labelled red) incorporated WT cells (shown in green) in two different ways. PA4782 and WT displayed a close to normal aggregate phenotype (as determined by average aggregate volume) however, aggregates were not clonal and contained a mixture of cells from both strains. PA2111 retained an entropic phenotype but was able to integrate WT cells into its distinct ‘raft-like’ formation. Mutant PA1864 alone is characterized as an impaired aggregator, however the addition of WT cells produced aggregates similar in volume to WT. Similar interactions were observed for all mutants. These preliminary findings of the WT able to influence the spatial organization of a mutant implies the occurrence of cooperative interactions during growth. Interestingly, during co-culture, entropic aggregates (PA4782 and PA2111) behave differently from each other. This raises many interesting questions about the combination of biological systems required for aggregation, specifically how the disruption of genes impacts physiology that relates to interactions with other cells. On a broader scale, the evolving accessibility of *Pa* cells within different formations to secreted factors and contact-dependent interactions may significantly influence the overall functionality of an aggregate.

## Conclusions

We provide evidence that amino acid uptake and metabolism, PQS, iron scavenging, oxidative stress response and LPS modification play fundamental roles in the process of *Pa* aggregation. The absence of WT aggregation and existence of multiple spatial phenotypes when these systems are disrupted supports this. Importantly, these mutations are not lethal and there is still much to be learned about aggregate metabolism, associated pathways, and their contribution to aggregation. The ability to temporally regulate aggregate genes is critical, where genes and operons need to be expressed at a particular time or growth phase to facilitate later functions (5102, 1864) that are required for WT aggregation. This result is especially important when considering using aggregation genes as potential therapeutic targets. At large, although are results relate directly to a model of chronic infection, we believe these data enhance our understanding of aggregate biology. More specifically unique requirements for aggregation in a CF-like environment could be used to inform better applications of existing therapeutics or the development of new ones against other aggregate forming bacteria.

## Acknowledgements

S.E.D is funded by the Cystic Fibrosis Foundation (CFF) (DARCH19G0, DARCH22P0) and start-up funds provided by the Department of Molecular Medicine, University of South Florida.

A.D.G was funded by a CFF student trainee award (GANNON22H0). Transposon mutants were provided by the University of Washington, funded by the CFF (SINGH19R0 and SINGH24R0). We thank Jessica Burns for guidance on R data analysis and Dr Lindsey Shaw and Lab (USF) for support analyzing RNA-seq data and discussion. We thank other members of the Darch Lab for discussion and reading of the manuscript.

## Author Contributions

*Conceived and designed the experiments*: S.E.D, A.D.G. *Performed the experiments*: A.D.G.

*Analyzed the data*: S.E.D, A.D.G and J.M *Wrote the paper*: S.E.D, A.D.G and J.M

## Data Availability

The RNA-seq data utilized in this manuscript for *Pseudomonas aeruginosa* will be submitted to NCBI. All other data sets generated for this study are included in the article or supplemental material.

## Materials and methods

### Strains, media and culture techniques

*Pseudomonas aeruginosa* PAO1 wildtype::*gfp* (containing plasmid pMRP9-1^1^ was cultured in standard lab media (LB) from frozen stock overnight at 37°C with shaking (200 rpm). Cells were back diluted 1:20 in fresh media and grown to log-phase ∼3 hours, then washed with PBS (pH7.0) before inoculation. *Pa* cells were inoculated at approximately 10^5^ cells per mL and incubated without shaking at 37°C as previously described^2^. Transposon mutants were obtained from the University of Washington and confirmed by Illumina sequencing (47). Strains were grown in LB (Gm50) and stocked at -80°C. We transformed Tn mutants were transformed with the fluorescent protein expressing plasmid pMP7605::*mCherry* (48). SCFM2 +mucin (250 mg) media was prepared as previously described (49). SCFM2 –mucin media was prepared identically with the exclusion of mucin. LB +mucin was prepared by adding mucin to sterile LB at the same concentration as SCFM2 +mucin media. LB –mucin consisted of sterile LB only.

### Comparison of aggregate and planktonic cultures and RNA extraction

After a period of 2, 4 or 8 hours, cultures were pelleted and re-suspended in DNA/RNA Shield (Zymo Research). Samples were flash frozen and stored at -80°C. The 0-hour timepoint was inoculated at a calculated OD_600_ equivalent of 8 hours growth for both LB and SCFM2 media, 0.53 and 1, respectively. This acted as a density control. After inoculation samples were grown statically at 37°C for 15 minutes and then harvested and frozen identically to other time points. Cell pellets were thawed on ice and 100μl of RNase/DNase free water added. Samples were transferred to bead beater tubes and 600μl of Qiagen lysis buffer added. Samples were bead beat at RT for 5 min, and then RNA extracted using the Qiagen RNeasy Mini kit and eluted in molecular grade water. RNA was stored at -80°C, and quantity and quality were measured using Qubit RNA High Sensitivity and RNA IQ kits (Thermofisher). Multiple technical replicates were preformed and pooled until there was at least 4 μg of total RNA for each biological replicate. To note, RNA was not recoverable from SCFM2 –mucin samples before 8 hours due to a lag phase in these conditions.

### RNA sequencing (RNA-seq) and differential expression (DE) analysis

Illumina RNA sequencing was performed by Novogene. FastQC was used to generate quality reports of the raw RNAseq FastQ reads (50). This data was used in MultiQC to make one comprehensive quality control report (51). Qiagen CLC Workbench was used for differential expression analysis. FastQ files generated by Novogene were imported into CLC Workbench as paired end reads with the following settings: minimum distance = 1, maximum distance = 1,000, Illumina options = remove failed reads, join reads from different lanes. The reference annotated gene track was created using reference genome PAO1_187 (NCBI) and reads were mapped using default mapping options. Reads were also mapped to annotated rRNA using default options; these mapped were excluded from differential expression analysis. The remaining unmapped reads were run through the CLC Workbench Differential Gene Expression Analysis workflow. For DE comparisons between time points within a single condition, the following settings were used: analysis was run in Batch, define batch units using metadata column: “Condition”, test differential expression due to: “Timepoint”, select “All group pairs for the Comparisons”. For DE comparisons between different conditions at one time point, the following options were used: analysis was run in Batch, define batch units using metadata column: “Timepoint”, test differential expression due to: “Condition”, select “All group pairs for the Comparisons”. Resulting tables of differentially expressed genes were filtered by fold change ≥ 2 and False Discovery Rate (FDR) P-value ≦ 0.05.

### Assessment of P. aeruginosa Tn-mutant aggregation

Transposon mutants were obtained from the PAO1 ordered transposon mutant library (University of Washington^3^). Each mutant was transformed with the fluorescent protein expressing plasmid pMP7605::*mCherry* (48). Mutants were cultured as described above and inoculated into 500μl SCFM2 +mucin at 37°C. For mixed strain experiments, WT PAO1 and mutants were inoculated 1:1. All images were acquired with the Zeiss LSM 880 confocal laser scanning microscope utilizing Zen image capture software. Bacterial cells were visualized via mCherry (excitation wavelength of 587 nm/emission wavelength of 610 nm) or GFP (excitation wavelength of 489 nm/emission wavelength of 508 nm) with a 63X oil immersion objective.

SCFM2 images were acquired by producing 512-by 512-pixel (0.26-by 0.26-μm pixels) 8-bit z-stack images that were 60 μm from the base of the coverslip. The total volume of the 60μm z-stack images were 1,093.5μm^3^. Control images of uninoculated SCFM2 were acquired by using identical settings to determine the background fluorescence for image analysis. Microscopy files were exported in CZI format.

### Image analysis

Image analysis. All imaging was performed with identical image capture settings. To determine the background fluorescence in SCFM2 as previously described (1). For aggregate studies in SCFM2, isosurfaces were produced for all remaining voxels after background subtraction with the surpass module in Imaris as previously described^2^. All image data were exported as a CSV and imported into R. Here, detected aggregate isosurfaces were then ordered by volume.

Objects that were >0.5 and <5.0 μm^3^ were categorized as dispersed biomass, and objects that were >5.0 μm^3^ were categorized as aggregated biomass. The total dispersed biomass was calculated as the sum of all the dispersed objects. Graphs were generated with GraphPad Prism 8.

### Computational Modeling

Since many pathway mapping programs struggle to accommodate hypothetical proteins, we used a predictive modeling pipeline to estimate the functions and pathways of these critically important genes (Fig. 2a). Protein sequences of identified hypothetical proteins were retrieved from the Pseudomonas Genome Database (52). Sequence homology to other proteins was determined using HMMER (53) .Structural homology was queried using both Dali (https://ekhidna2.biocenter.helsinki.fi/dali/) and SWISS-Model (54) .This structural homology information was used to predict protein structures using AlphaFold2 ColabFold v1.5.2 (55).

Protein-protein interactions were mapped using STRING pathway analysis, which utilizes information from experiments, as well as large omics datasets (56). This information taken in conjunction with the predicted function allowed us to postulate mechanisms and pathways that these unannotated proteins are likely to be involved in (Table 1).

## Supplementary figure legends

Table S1 – The 50 most dysregulated differentially expressed genes from aggregate samples. 3 biological replicates, significance cutoffs 2-fold change, FDR 0.05.

Table S2 – Differentially expressed pyocins in aggregates. Highlighted pyocins are found in both mucin-containing conditions.

Figure S1 – Genes specific to CF-like environment are needed for WT aggregation. (a) Venn diagram showing number of unique and overlapping genes between several *in vitro* models and *ex vivo* samples. (b) Total biomass over time of transposon mutants grown in SCFM2. Entropic mutants maintain large populations with rapid doubling times, while impaired mutants have a near or slower than WT growth rate. 3 biological replicates +/-SEM.

Figure S2 – Prediction pipeline. This pipeline uses sequential and structural homology as a basis to predict the function and pathway of hypothetical proteins important in aggregates. AlphaFold is used to model protein complexes, and STRING is used to predict interactions with other proteins.

Figure S3 – WT co-culture. (a) Comparisons of mutant and WT total biomass, average aggregate size, and number of planktonic cells at 5 and 10 hours. (b) Comparisons of WT total biomass, average aggregate size, and number of planktonic cells at 10 hours. Each WT was co-cultured with the respective mutant, resulting in significant differences in several growth measures. 3 biological replicates +/-SEM, significance of mutants vs. WT calculated using ordinary two-way ANOVA (P value <0.0001) with Fischer’s LSD multiple comparisons test (alpha 0.05). WT-WT comparisons used Kruskal-Wallis test (P value 0.0051) and Dunn’s multiple comparisons test (alpha 0.05).

